# First isolation of virulent *Tenacibaculum maritimum* strains from diseased orbicular batfish (*Platax orbicularis*) farmed in Tahiti Island

**DOI:** 10.1101/2021.03.15.435441

**Authors:** Pierre Lopez, Denis Saulnier, Shital Swarup-Gaucher, Rarahu David, Christophe Lau, Revahere Taputuarai, Corinne Belliard, Caline Basset, Victor Labrune, Arnaud Marie, Jean François Bernardet, Eric Duchaud

## Abstract

The orbicular batfish (*Platax orbicularis*), also called ‘Paraha peue’ in Tahitian, is the most important marine fish species reared in French Polynesia. Sudden and widespread outbreaks of severe ‘white-patch disease’ have occurred since 2011 in batfish farms one to three weeks after the transfer of juveniles from bio-secured hatcheries to lagoon cages. With cumulative mortality ranging from 20 to 90%, the sustainability of aquaculture of this species is severely threatened.

In this study, we describe for the first time the isolation from diseased batfish of several strains belonging to the species *Tenacibaculum maritimum*, a major pathogen of many marine fish species. Histopathological analysis, an experimental bath challenge and a field monitoring study showed that *T. maritimum* is associated with white-patch disease. Moreover, molecular and serological analyses performed on representative strains revealed some degree of genetic diversity among the isolates, a finding of primary importance for epidemiological studies and for the development of management and control strategies such as vaccination.

## 1 Introduction

The orbicular batfish (*Platax orbicularis*, family *Ephippidae*) is a fish species inhabiting the top 30 metres depth over tropical reefs of the western Pacific [1]. It is widely distributed from the Indo-Pacific region to the Red Sea in East Africa, with a northern limit in south Japan and a southern limit in Australia and New Caledonia [2]. In French Polynesia, the batfish is highly appreciated as a food by the local population, but the wild stock is rapidly decreasing, probably as a result of high fishing pressure and climate change that severely disturb the coral communities [3]. To promote local sustainable aquaculture, the governmental department of marine resources (DRM) decided to focus effort on captive breeding and rearing of orbicular batfish. Since 2011, the governmental hatchery VAIA (Vairao, Tahiti, French Polynesia) has produced eighty thousand fry annually, reared inland in bio-secured conditions (i.e. in filtered and UV treated seawater). Four production cycles are completed per year, allowing an annual production of 50–80 metric tons. One to two-month-old juvenile fish (mean weight, 1 and 10 g) are then transferred to different fish farms in lagoons. However, recurrent mortalities occur almost systematically during the first two months of growth in the lagoon cages, causing losses of 20–90% of production and threatening the sustainability of aquaculture based on this species. One to three weeks following their transfer to net cages in the lagoon, the fish show symptoms of disease, with loss of appetite, frayed fins, whitish patches on the tegument, followed by ulcers, necrosis and death. Little was known about the status of pathogens and diseases associated with this tropical fish species under local farming conditions and, consequently, this disease was simply named ‘white-patch disease’, based on the clinical symptoms. Light microscopy examination of fragments of skin lesions revealed abundant rod-shaped and gliding bacteria potentially belonging to the genus *Tenacibaculum*.

The genus *Tenacibaculum* (family *Flavobacteriaceae*, phylum *Bacteroidetes*) currently comprises 31 validly named species (http://www.bacterio.net/tenacibaculum.html), all retrieved from marine environments [4]. Among these, *T. dicentrarchi, T. discolor, T. finnmarkense, T. gallaicum, T. maritimum, T. piscium* and *T. soleae* are responsible for ulcerative conditions that affect a large variety of cultured and wild marine fish species and are collectively known as tenacibaculosis [5]. Tenacibaculosis is generally associated with gross external lesions such as ulcerative and necrotic skin lesions, haemorrhagic mouth, frayed fins and tail rot [5]. The disease was originally described in 1977 in cultured red (*Pagrus major*) and black (*Acanthopagrus schlegeli*) seabream in Japan [6]. The causative agent was subsequently identified as *Tenacibaculum maritimum* (formerly *Flexibacter maritimus*) [7], a filamentous, Gram negative, gliding bacterium. Since then, *T. maritimum* has been shown to be responsible for considerable losses in marine aquaculture worldwide, affecting a large variety of wild and cultured marine fish species. For example, *T. maritimum* has been found to be associated with mortality events occurring in Atlantic salmon (*Salmo salar*) in Australia [8], Chinook salmon (*Oncorhynchus tshawytscha*) in Canada [9], rainbow trout (*Oncorhynchus mykiss*) in Australia [8], sole (*Solea senegalensis*) and turbot (*Scophthalmus maximus*) in Spain [10], sea bass (*Dicentrarchus labrax*) in Europe [11], Japanese flounder (*Paralichthys olivaceus*) in Japan [12], and black damselfish (*Neoglyphieodon melas*) and Picasso triggerfish (*Rhinecanthus assasi*) in Egypt [13]. Although other *T. maritimum* isolates have also been retrieved from outbreaks in other countries and host fish species, pathogenicity has not been confirmed using experimental challenges in any of these examples [14,15].

The aims of the present study were to investigate the recurrent and acute outbreaks occurring in *Platax orbicularis* farms in French Polynesia, to characterize the causative agent using bacteriological, histological and molecular analyses and then to conduct experimental challenges to confirm this causality, thus fulfilling Koch’s postulates.

## 2 Materials and methods

### 2.1 Ethic statement

In the absence of *adhoc* ethical committees in French Polynesia, *in vivo* experiments reported in the present study fulfill all the sections of deliberation no 2001-16 APF from the Assembly of French Polynesia issued in the Journal Officiel de Polynésie française on the 1st February 2001, dealing on domestic or wild animal welfare. Nevertheless, we used several guidelines in the present study and followed animal care and ethic guidelines (16,17). In particular, fish were euthanized using an overdose of Benzocaine (150 mg L^-1^ stock solution prepared in ethanol). This method of euthanasia, reproducible and safe to the operator, induces a depression of the central nervous system activity, rapid unconsciousness and death of *P. orbicularis*, without compromising further microbiological and histological analyses. The criterion used to exercise humane endpoint was that moribund fish displaying the typical sign of white-patch disease had lost their ability to maintain an upright position in the tanks and were not evasive to netting. During all experiments of this study, fish were monitored by trained fish health specialists to make sure that the ethical aspects were secured.

### 2.2 Sampling of diseased fish

Ten symptomatic orbicular batfish (mean weight 5.1 +/- 2.3 g) were recovered from two different farms located in Tahiti island. They were sampled during two severe outbreaks in 2013 and 2016 that had caused >50% cumulative mortality just 2 weeks after the fish had been transferred from the VAIA hatchery to net cages in the Tahiti lagoon. All fish showed erosion and ulceration of the skin surface. After being euthanized with an overdose of Benzocaine (150 mg L^-1^), they were examined by microscopy and microbiological and histological techniques.

### 2.3 Direct microscopic examination and isolation of bacteria

Skin lesion scrapings from moribund batfish were collected using sterile surgical scalpels. Wet mount preparations were then examined under a light microscope (Leica DM 1000 LED). For each bacterial isolation, a sterile swab cotton-tipped applicator (COPAN) was used. Smears of skin samples were deposited onto plates of *Flavobacteriaceae*-selective marine agar (FSMA) developed by an accredited veterinary diagnostic laboratory (Labofarm, Loudéac, France). A total of ten dominant bacterial strains were isolated after sub-culture (Table 1).

**Table 1.** List of bacterial strains retrieved from farmed *Platax orbicularis* affected by white-patch disease, with their sources and characteristics. Virulence was evaluated by experimental bath challenge on *T. maritimum-free* batfish. Significant differences (p<0.05) in mortality rate (see paragraph 3-3) between non-infected and infected fish are indicated by ‘yes’ or ‘no’. Isolate identification was performed using EzBioCloud software [18], based on >99% identity of their 16S rRNA sequences with the closest type strain. ST refers to the MLST sequence type. n/a: not analysed.

| Strains | Source and date of isolation | GPS location | 16S rDNA<br>GenBank accession<br>number | Bacterial species | Virulence | ST |
| --- | --- | --- | --- | --- | --- | --- |
| TFA4 | Skin lesions, Tautira lagoon, Tahiti, 2013 | 17°47'50'' S,<br>149°07'14'' W | MW690171 | *T. maritimum* | yes | ST168 |
| Aq 9-66 | Skin lesions, Tautira lagoon, Tahiti, 2013 | 17°47'50'' S,<br>149°07'14'' W | MW690177 | *T. mesophilum* | no |  |
| Aq 9-67 | Skin lesions, Tautira lagoon, Tahiti, 2013 | 17°47'50'' S,<br>149°07'14'' W | MW690178 | *T. mesophilum* | no |  |
| Aq 16-83 | Skin lesions, Vairao lagoon, Tahiti, 2016 | 17°48'22'' S,<br>149°17'36'' W | MW690172 | *T. maritimum* | n/a | n/a |
| Aq 16-84 | Skin lesions, Vairao lagoon, Tahiti, 2016 | 17°48'22'' S,<br>149°17'36'' W | MW690173 | *T. maritimum* | yes | n/a |
| Aq 16-85 | Skin lesions, Vairao lagoon, Tahiti, 2016 | 17°48'22'' S, 149°17'36'' W | MW690174 | *T. maritimum* | n/a | ST167 |
| Aq 16-87 | Skin lesions, Vairao lagoon, Tahiti, 2016 | 17°48'22'' S, 149°17'36'' W | MW690175 | *T. maritimum* | yes | n/a |
| Aq 16-88 | Skin lesions, Vairao lagoon, Tahiti, 2016 | 17°48'22'' S, 149°17'36'' W | MW690176 | *T. maritimum* | n/a | ST167 |
| Aq 16-89 | Skin lesions, Vairao lagoon, Tahiti, 2016 | 17°48'22'' S, 149°17'36'' W | MW690180 | *T. maritimum* | n/a | ST167 |
| Aq 16-91 | Skin lesions, Vairao lagoon, Tahiti, 2016 | 17°48'22'' S, 149°17'36'' W | MW690179 | *T. mesophilum* | n/a |  |

### 2.4 Histopathological examination

Skin fragments of approximately 1 cm^2^ were collected from four fish showing typical skin lesions of the white-patch disease using a sterile surgical scalpel. Samples were fixed in Davidson’s solution (25% formaldehyde, 37.5% ethanol, 12.5% acetic acid and 25% water) for 48 hours at room temperature, then washed and kept in 70% ethanol. Skin samples were progressively dehydrated in an ascending series of alcohol (70 to 100% ethanol) followed by a xylene bath, using a dehydration automate (Leica, ASP 300S), then embedded in paraffin, cut into 3-μm sections using a rotary microtome (Microm HM 340E, Thermo Fisher Scientific) and stained with haematoxylin-eosin (H-E) using a fully-automated integrated stainer (Leica, CV5030 autostainer XL). Several sections were analysed to ensure reproducibility, using a Leica DM 1000 LED microscope equipped with a Dino-Lite camera (AnMo Electronics).

### 2.5 Molecular and serological studies

16S rDNA sequences were PCR-amplified using the universal 27F and 1492R primers (Table S1) and the purity and length of the amplicons were verified by agarose gel electrophoresis. Amplicons were Sanger sequenced by GATC-biotech (https://www.gatc-biotech.com) using the six universal sequencing primers listed in Table S1. For each strain, the six sequences were visualized and aligned to create a consensus sequence (with > 2X coverage over 80% of the sequences) using Benchling software (2020). For primary taxonomic assignation, the 16S rRNA consensus sequences were searched against the EzBioCloud database [18] (accession numbers are given in Table S2). In addition, a tentative phylogenetic tree was constructed using the MAFFT online service [19]. The evolutionary distance was calculated using 1000 bootstrap replicates (Fig S1).

To characterize the genetic diversity of presumptive *T. maritimum* strains in greater depth, multi-locus sequence analysis (MLSA) was performed on four selected isolates (TFA4, Aq 16-85, Aq 16-88, Aq 16-89) using sequences retrieved from their draft genomes [20]. These isolates were selected according to their background information: Aq 16-85, Aq 16-88 and Aq 16-89 were sampled from three different infected fish during an outbreak at the Vairao fish farm in 2016, while strain TFA4 was isolated from a symptomatic fish at the Tautira fish farm in 2013 (Table 1). The MLSA profile defined by Habib et al. (2014) [21] consists of the sequences of seven housekeeping genes (*atpA, gyrB, dnaK, glyA, infB, rlmN* and *tgt*). The profiles of the new allele and sequence types (ATs and STs, respectively), were generated and analysed using the *Tenacibaculum* pubMLST database (https://pubmlst.org/tenacibaculum/) [22]. Results were visualized using the incremented Interactive Tree of Life (iTOL) v3 tool [23].

To examine the isolates identified as *T. maritimum* more closely, their serotype was determined as described by Avendaño-Herrera et al. (2004b) [24]. This method uses a slide agglutination test and a dot blot assay on both whole-cell preparations and heat stable O antigens of each strain. Antisera against serovars O1 (PC503.1), O2 (PC424.1) and O3 (ACC13.1) were used in all assays.

### 2.6 Experimental infection by immersion and quantification of *Tenacibaculum maritimum* in mucus samples by real-time qPCR

A batch of *Platax orbicularis* fingerlings (mean weight 9.7 +/- 2.6 g) reared at the VAIA bio-secured hatchery was transferred for acclimatization to a 1-m^3^ fibreglass tank containing seawater (salinity, 32 PPT; water temperature of 26–27°C) for 10 days. Prior to infection, 494 fish were randomly selected and transferred to nine 150-L tanks (50 fish per tank) filled with 5-μm filtered seawater. Three groups were tested in triplicate: (i) non-infected fish (NIF), (ii) fish infected with *T. maritimum* strain TFA4 (IF) and (iii) fish with impaired mucus (IM) infected with *T. maritimum* strain TFA4 (IM-IF). The mucus of the latter fish was partially removed by gently wiping one side of the fish with a sponge soaked in filtered seawater. Fish were challenged with a pure culture of strain TFA4 obtained by incubation at 27°C for 48 h (stationary phase) in autoclaved nutrient broth composed of 4 g L^-1^ peptone and 1 g L^-1^ yeast extract (Becton, Dickinson and Co., Sparks, MD, USA) in 5μm-filtered seawater under orbital shaking at 200 rpm. Bath challenges were performed for two hours with strain TFA4 at a final concentration of 5.3 10^4^ CFU mL^-1^ for groups IF and IM-IF or with nutrient broth in the case of the mock-treated NIF control group. The infected fish were then rinsed twice with filtered seawater to remove all non-adherent bacteria, and fish from the NIF control group were manipulated in the same way. Mortality was monitored twice daily from day 0 (D0) until day 5 (D5) at which point the fish were euthanized using 150 mg L^-1^ Benzocaine. Any fish that died or were found moribund over the experimental period (D0-D5) were promptly removed from the tanks during the monitoring. The non-parametric Kaplan–Meier method (R package *survival*) was used to test for differential survival performances among groups at the same date or within groups throughout the sampling period. Differences were considered significant at *P* < 0.05.

At 24 h post-infection, before the onset of mortality, four fish displaying skin lesions were randomly sampled from the two infected groups (IF and IM-IF) in addition to four fish from the NIF group. These were used to quantify *T. maritimum* cells in fish mucus using TAQMAN real time PCR (see primers and probe in suppl. Table 1) following the protocol developed by Fringuelli et al. (2012) [23] with minor modifications. Briefly, skin mucus samples were obtained from lesions on symptomatic fish using cotton swabs (COPAN), directly diluted in 1.5-mL micro-centrifuge tubes containing 0.5 mL of lysis solution (0.1 M EDTA pH 8; 1 % SDS and 200 μg ml^-1^ proteinase K) and incubated overnight at 55°C. DNA was extracted using the conventional phenol/chloroform/isoamyl alcohol (25/24/1) method. DNA quantity and purity were assessed using a NanoDrop ND 1000 spectrophotometer (Thermo Fisher Scientific). In order to obtain a standard curve, bacterial cells of strain TFA4 from a stationary phase culture in nutrient broth were enumerated using a Malassez counting chamber (2.35 10^8^ bacteria ml^-1^) and DNA from 1 ml of the bacterial suspension was extracted. The DNA was then spiked at a final concentration of 1.33 ng μl^-1^ in salmon sperm gDNA (SSD, Thermo Fisher) at 10 ng μl^-1^ in artificial seawater (ASW, Sigma), then serially diluted 10-fold in SSD at 10 ng μl^-1^ in ASW. A linear range of values was obtained for PCR amplification on a Mx3000 Thermocycler (Agilent) using Brilliant III Ultra-Fast QPCR Master Mix (Agilent) following the supplier’s recommendations (5 μl DNA at 10 ng μl^-1^ in a total reaction volume of 20 μl), with six successive sample 10-fold dilutions tested in triplicate. Cycle threshold (Ct) values ranged from 16.05 to 33.14, corresponding to 1.44 10^5^ to 2.06 10^1^ cells of strain TFA4 per PCR well, while correlation (linear regression with r^2^ coefficient) and qPCR reaction efficacy were 0.995 and 99.9%, respectively.

### 2.7 Detection and quantification of *Tenacibaculum maritimum* during a field episode of tenacibaculosis

Juvenile batfish (mean weight 10 +/- 3.3 g) reared in the bio-secured facilities of the VAIA hatchery were carefully transferred (D0) to the Tahiti Fish Aquaculture farm in Tautira lagoon and kept in a single net cage of 1 m^3^ (167 fish/m^3^ density). Ten fish were collected at five sampling times: day 6 before transfer (D-6) (i.e. in the VAIA hatchery) and D1, D9, D17 and D36 post-transfer to Tautira lagoon. When gross signs of the white-patch disease were observed, five moribund fish (symptomatic) and five apparently healthy ones (asymptomatic) were sampled. These fish were euthanized as detailed above and the liver, posterior intestine and some skin mucus (collected with a cotton swab in the lesion area in the case of symptomatic fish) were individually and aseptically sampled and preserved in 500 μl RNAlater (Ambion) at −80°C. Approximatively 100 mg of tissue were used to quantify *T. maritimum* by qPCR [23]. Throughout this survey, no curative treatments were given, mortality was monitored daily and moribund animals (euthanatized with an overdose of benzocaine) or dead fish were removed and discarded.

## 3 Results

### 3.1 Microscopic examination and isolation of bacteria

Two severe white-patch disease outbreaks, occurring within the first 2 months following transfer to net cages in a lagoon, were recorded in 2013 and 2016. These outbreaks occurred in two geographically distinct fish farming areas of Tahiti island (Tautira lagoon: 17°47’50” S, 149°07’14” W, and Vairao lagoon: 17°48’22” S, 149°17’36” W) with cumulative mortality reaching 80% and 62%, respectively. The main clinical signs were loss of appetite, erratic swimming and ulcerative skin lesions (Fig 1A). Wet mount examination of the skin mucus of diseased batfish revealed a significant amount of long (6.3 +/- 0.6 μm) and rod-shaped bacteria (Fig 1B).

**Fig 1.**
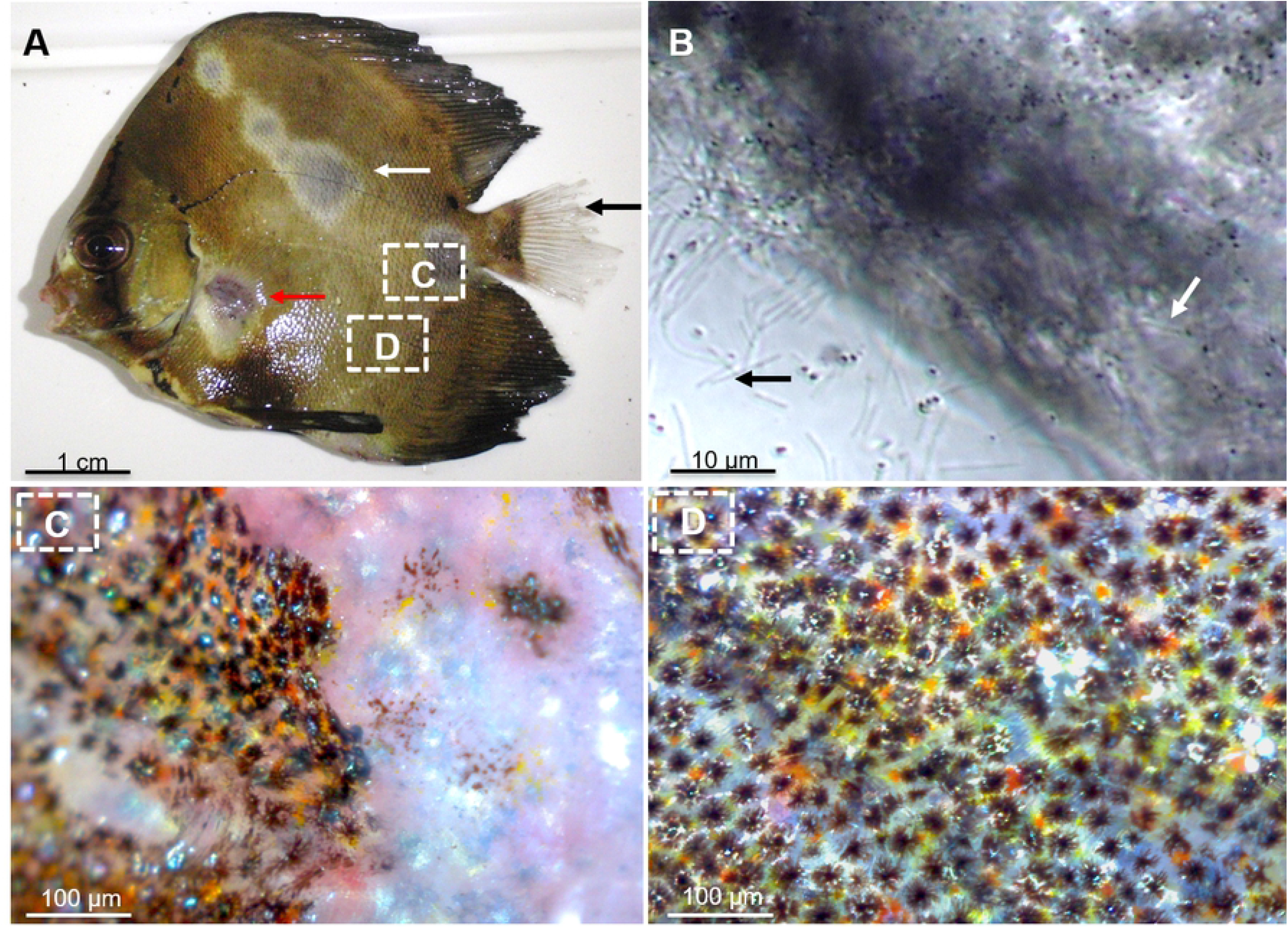
Examination of fish lesions. A) Gross clinical signs of the white-patch disease of batfish characterized by: (i) circular discoloration areas of various sizes, apparently randomly distributed on the skin surface; (ii) skin lesions, ulcers, scale loss (white arrow) and areas of haemorrhagic necrosis (red arrow); and (iii) frayed (usually caudal) fins (black arrow). B) Microscopic examination of skin lesions reveals abundant, long, slender, rod-shaped bacteria. Numerous bacteria remain adhered to the fish scales (white arrows) while others detached after the fragment of lesion was crushed. C) View of the skin surface at the interface between apparently healthy and damaged zones. D) An apparently healthy zone observed under a surgical Q-Scope microscope (AnMo Electronics).

Histopathological examination of skin lesions from moribund batfish revealed that the epidermis and dermis were severely damaged, with clusters of filamentous, *Tenacibaculum*-like bacteria and scattered inflammatory cells (Fig 2). In contrast, no evidence of histopathological changes was noticed in the internal organs.

**Fig 2.**
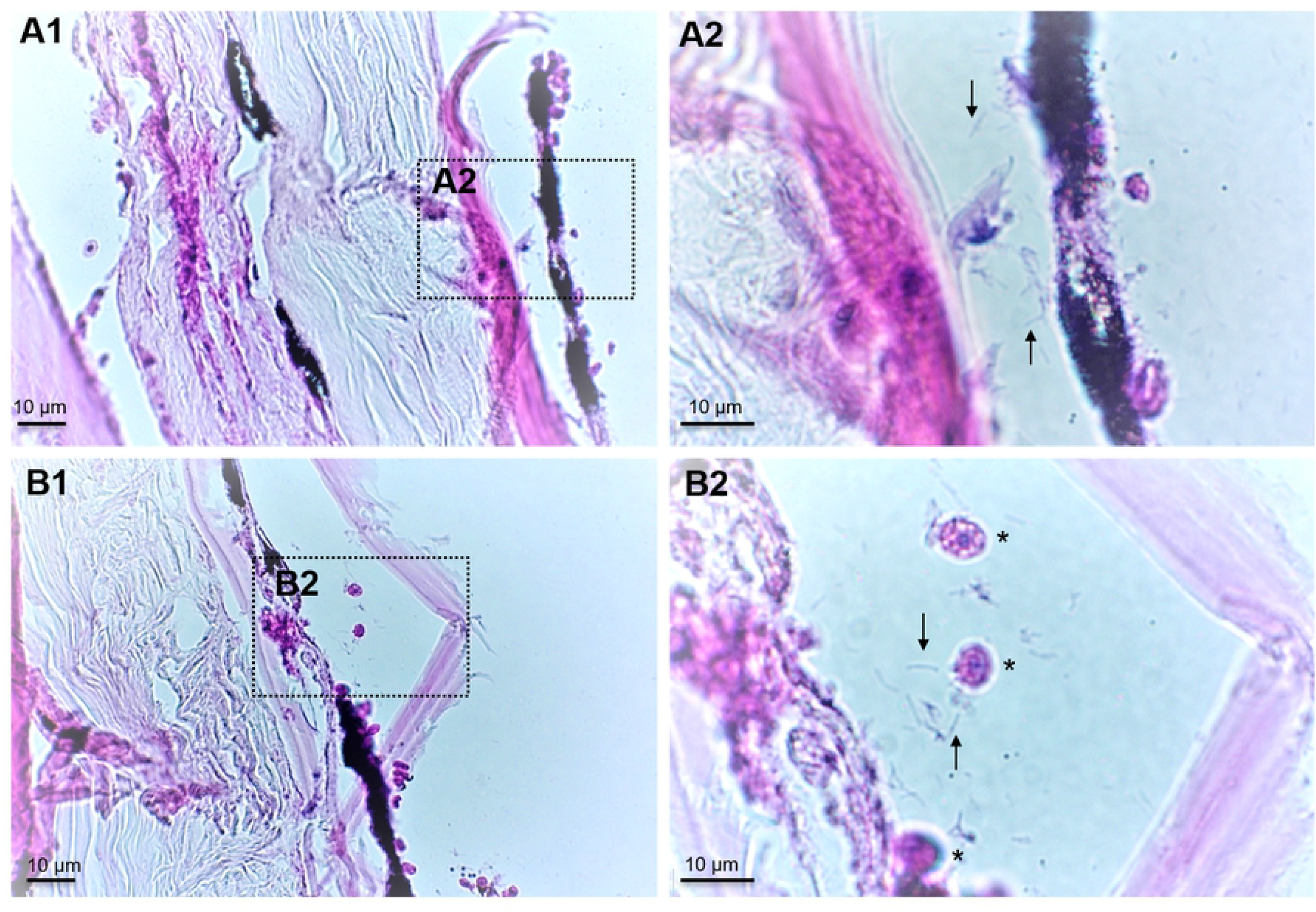
Two representative cross sections (H-E staining) of *Platax orbicularis* fingerlings affected by white-patch disease. A1 and B1: Severe necrosis affecting the hypodermis and dermis layers with invasion of *Tenacibaculum* cells (arrows) visible at a higher magnification (A2 and B2) and detection of inflammatory cells in damaged areas (asterisks).

Ten strains were isolated from samples of 10 moribund batfish exhibiting the typical signs of white-patch disease using *Flavobacteriaceae*-selective marine agar (Table 1). Two different colony morphotypes were observed after 48 h of incubation at 27°C: the first morphotype consisted of pale, translucent colonies with uneven edges, extremely adherent to the agar (Fig 3A), while the second morphotype consisted of orange, opaque, diffuse and strongly iridescent colonies (Fig 3B, C).

**Fig 3.**
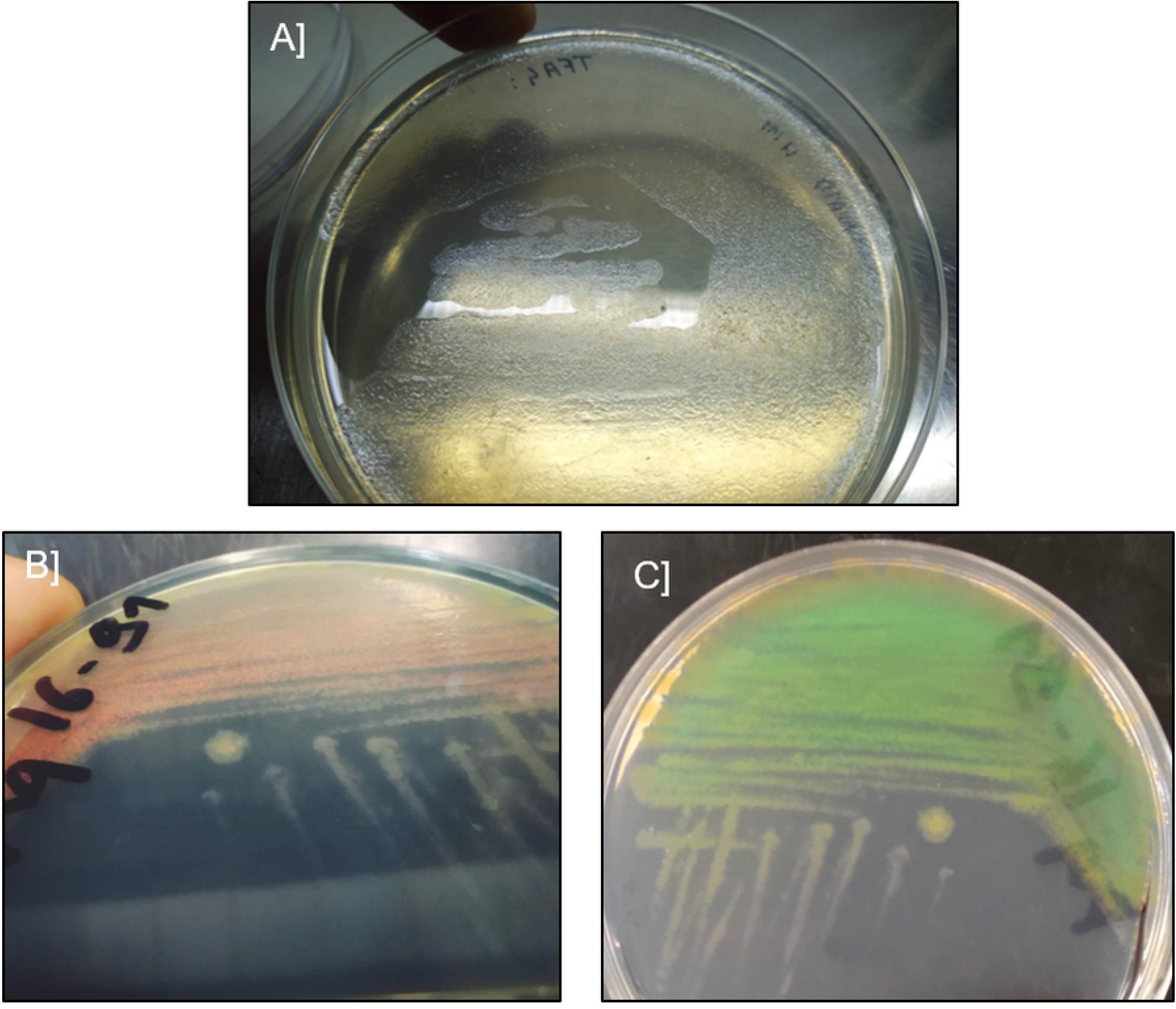
Representative strains isolated from symptomatic *Platax orbicularis*. A) colonies of *Tenacibaculum maritimum*, strain TFA4. B) and C) colonies of *Tenacibaculum mesophilum*, strain Aq 16-91, with different camera shooting angles revealing the iridescent phenotype.

### 3.2 Genomic and serological characterization

Analysis of nearly complete 16S rRNA sequences revealed that all isolates belonged to the genus *Tenacibaculum.* Seven strains (TFA4, Aq 16-83, Aq 16-84, Aq 16-85, Aq 16-87, Aq 16-88 and Aq 16-89) shared 99.65% to 99.79% sequence identity with the *T. maritimum* type strain NBRC 15946^T^, with at least 98,9% coverage. The three remaining strains (Aq 9-66, Aq 9-67 and Aq 16-91) displayed 99.37 to 99.93% sequence identity with the *T. mesophilum* type strain DSM 13764^T^, with at least 98,9% coverage. A tentative phylogenetic tree was drawn using MAFFT (Fig S1). The seven strains displaying the first morphotype clustered with the *T. maritimum* type strain, while the three strains belonging to the second morphotype clustered with the *T. mesophilum* type strain; bootstrap values were 100% and 91%, respectively.

The results of the MLSA analysis (Fig 4) performed on *T. maritimum* strains TFA4, Aq 16-85, Aq 16-88 and Aq 16-89 showed that none matched exactly with any of the sequence types (ST) already described in the pubMLST database. They were therefore treated as belonging to new STs: ST168, which was attributed to strain TFA4; and ST167, which was attributed to strains Aq 16-85, Aq 16-88 and Aq 16-89. Analysis of the number of locus variants revealed that these novel STs only share three allele types (AT), corresponding to loci *gyrB, infB* and *rlmN*, which reveal genetic heterogeneity among these two groups of isolates. The single and double locus variant analyses (SLV and DLV) were fairly congruent with the phylogenetic tree based on the concatenated nucleotide sequences of the seven housekeeping genes (Fig 4). SLV analysis showed that ST168 (TFA4) shares 6/7 loci with ST2, which up to now only included strain ACC13.1 (referenced as 002 in the pubMLST database), isolated from a diseased Senegalese sole (*Solea senegalensis*) in Portugal. In the DLV, TFA4 shared 5/7 loci with ST3, 4, 10, 35, 36, and 130, which essentially comprise isolates from the south of Europe (except for strain 4646, isolated in Australia). Interestingly, strains Aq 16-85, Aq 16-88 and Aq 16-89 (ST167) formed a singleton, meaning that they displayed at least three different ATs compared with all the strains included in the pubMLST database. These results revealed the existence of at least two genetically distinct groups of *T. maritimum* isolates in Tahitian fish farms.

**Fig 4.**
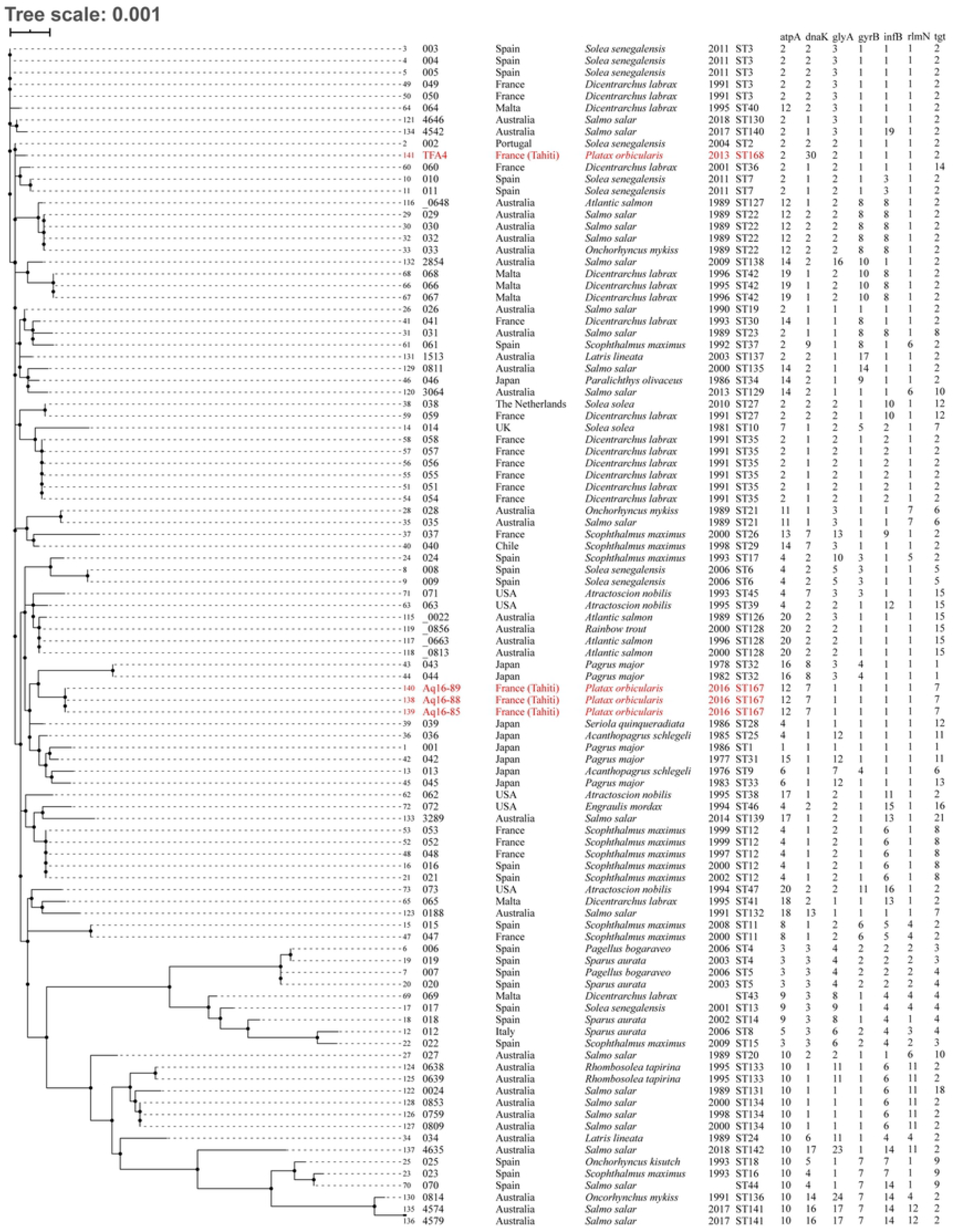
Genomic and background information on all *T. maritimum* isolates in the pubMLST database. Strains isolated in this study are shown in red type. Neighbor-joining tree based on the concatenated nucleotide sequences of the 7 housekeeping genes (3894 bp). Other information: strain number, strain name, country of isolation, fish host species, year of isolation, sequence type, and its allelic combination.

Serological analysis revealed two different serogroups among the *T. maritimum* strains. Slide agglutination tests showed that strain TFA4 specifically reacted with the anti-O3 antiserum, while strains Aq 16-85, Aq 16-88 and Aq 16-89 specifically reacted with the anti-O1 antiserum.

### 3.3 Pathogenicity assays using an immersion challenge

All batfish that were experimentally infected with *T. maritimum* TFA4 using the immersion challenge (IF group) exhibited typical clinical signs of white-patch disease starting from 24 h post-infection (PI), although no mortality was recorded at this time point (Fig 5). At 30 h PI, fish in this group underwent significant (chi-squared-test with simulated p-value correction, P = 0.017) mortality, with a survival probability of 94.9% compared with 100% (no death events) in the non-infected fish group (NIF). A sudden mortality event was observed in the IF group between 30 and 48 h PI, with 94.9% and 25% survival probabilities, respectively. From 82 h until the end of the monitoring period (120 h PI), no increase in mortality was observed (7.1% survival probability for the IF group at 72 h PI to 120 h PI) even though all batfish displayed typical clinical signs of the disease. Nevertheless, the intensity of ulcerative skin lesions (i.e. the number and area of whitish patches) in the IF group from 72 h to 120 h PI was lower than that observed before 72 h PI (data not shown). In the IM-IF group, not only did batfish experience the highest mortality rate because all fish died, giving a survival probability of 0% from 30h PI, but they also died significantly earlier than those of the IF group with intact skin mucus (log-rank test comparing the survival curves from 0 h PI to 30 h PI, P = 0.02). No mortality was recorded in the NIF control group during the entire trial.

**Fig 5.**
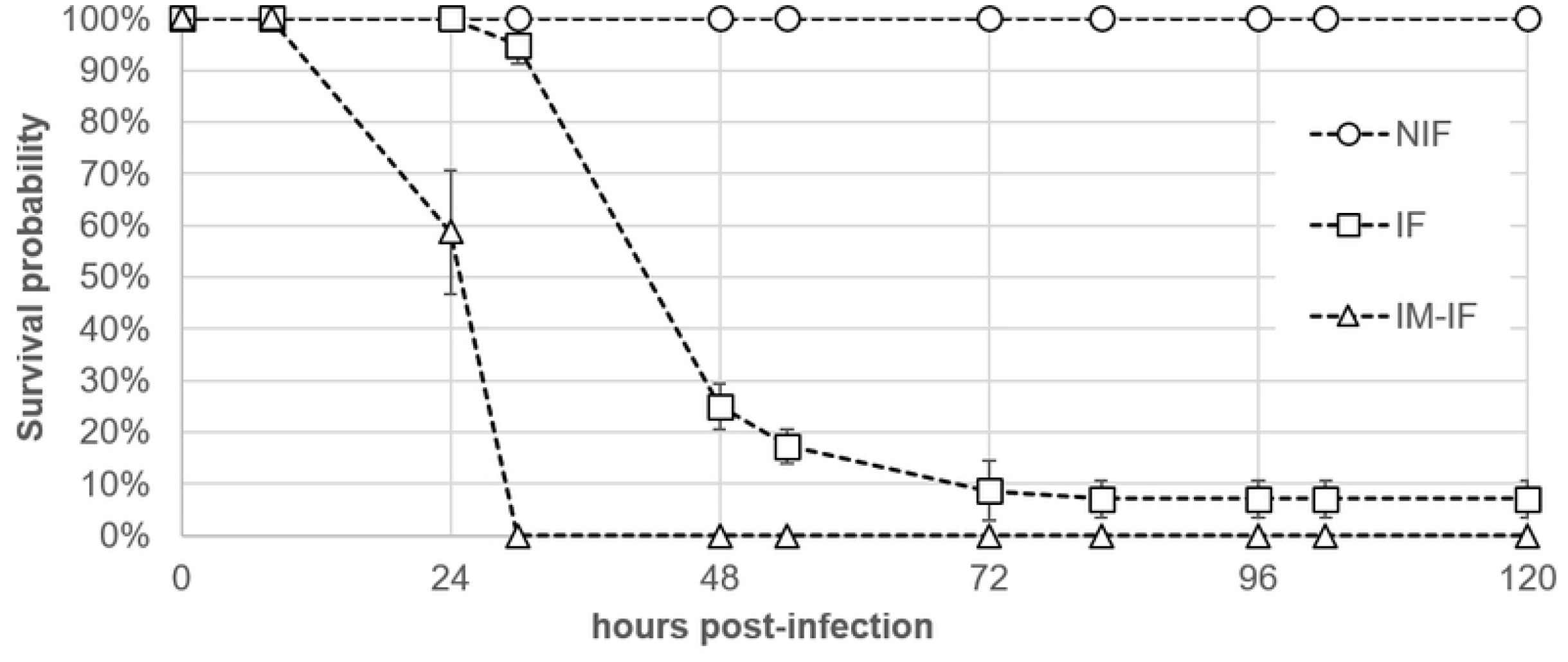
Survival curves of bath-challenged batfish *Platax orbicularis.* NIF, non-infected fish; IF, fish infected with 5.3 10^4^ CFU mL^-1^ of strain TFA4 for 2 hours; IM-IF, infected fish from which mucus had been partially removed before the bath challenge.

All skin samples that were collected from diseased fish in the IF group before the onset of mortality were found positive by real time PCR assay, with an average load of 7.8 10^8^ +/- 1.4 10^8^ *T. maritimum* bacteria per μg of total extracted DNA. In contrast, no *T. maritimum* was detected in any of the four analysed skin samples in the NIF group. The virulence potential of two (Aq 16-84 and Aq 16-87) and three (Aq 9-65, Aq 9-66 and Aq 9-67) strains belonging to the species *T. maritimum* and *T. mesophilum*, respectively, was also evaluated using the same immersion challenge protocol. Results of this separate trial showed that the two *T. maritimum* strains exhibited levels of virulence similar to that of strain TFA4, whereas the three *T. mesophilum* strains were avirulent at a similar infection dose (6.1 10^4^ CFU mL^-1^), with no mortality recorded in the groups IF and NIF during a seven-day post-infection survey (data not shown).

### 3.4 Kinetics of *Tenacibaculum maritimum* infection during a field episode of tenacibaculosis

To gain insight into *T. maritimum* pathogenesis under natural field conditions, a batch of batfish was monitored from its production under bio-secured conditions at the VAIA hatchery to its rearing in a net cage on a private farm in Tautira lagoon. Soon after the transfer to the net cage in the lagoon, a severe outbreak of white-patch disease was observed, with the first typical signs appearing from D1 post-transfer and mortality from D3 (Fig 6).

**Fig 6.**
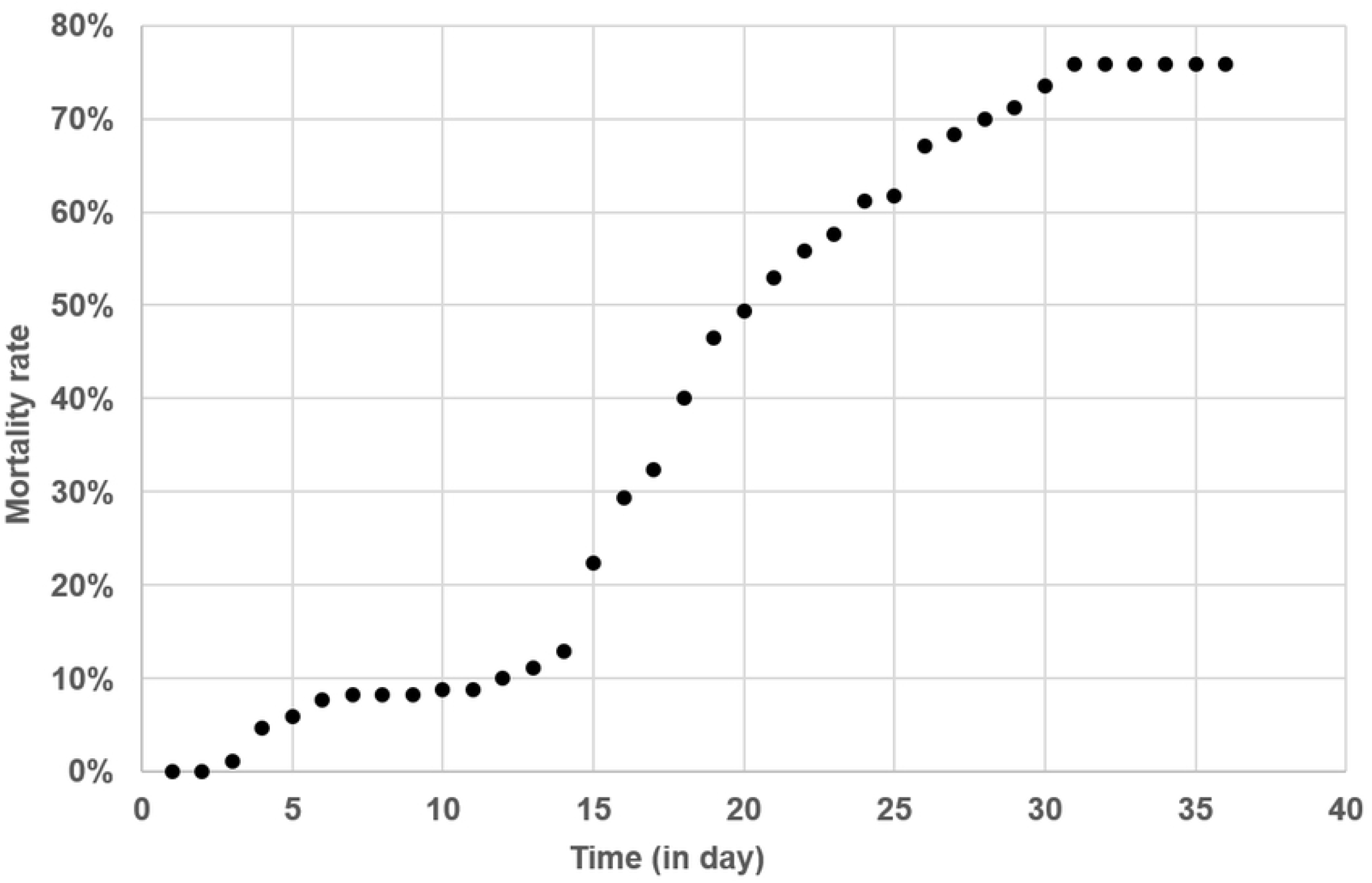
Cumulative mortalities of a batch of batfish during a natural outbreak following their transfer to a net cage in Tautira lagoon. Fish were 10 g (mean weight) and reared at an initial density (D0) of 167 fish/m^3^.

Two peaks of mortality occurred, at D3-D6 and D13-D31. The second peak was higher, with cumulative mortalities increasing significantly from 11.2% at D13 to 74.5% at D31. No subsequent mortality then occurred among the surviving batfish until the end of the study period (D36).

Six days before the transfer to the lagoon cage (D-6) all sampled fish were qPCR negative for *T. maritimum* (Fig 7).

**Fig 7.**
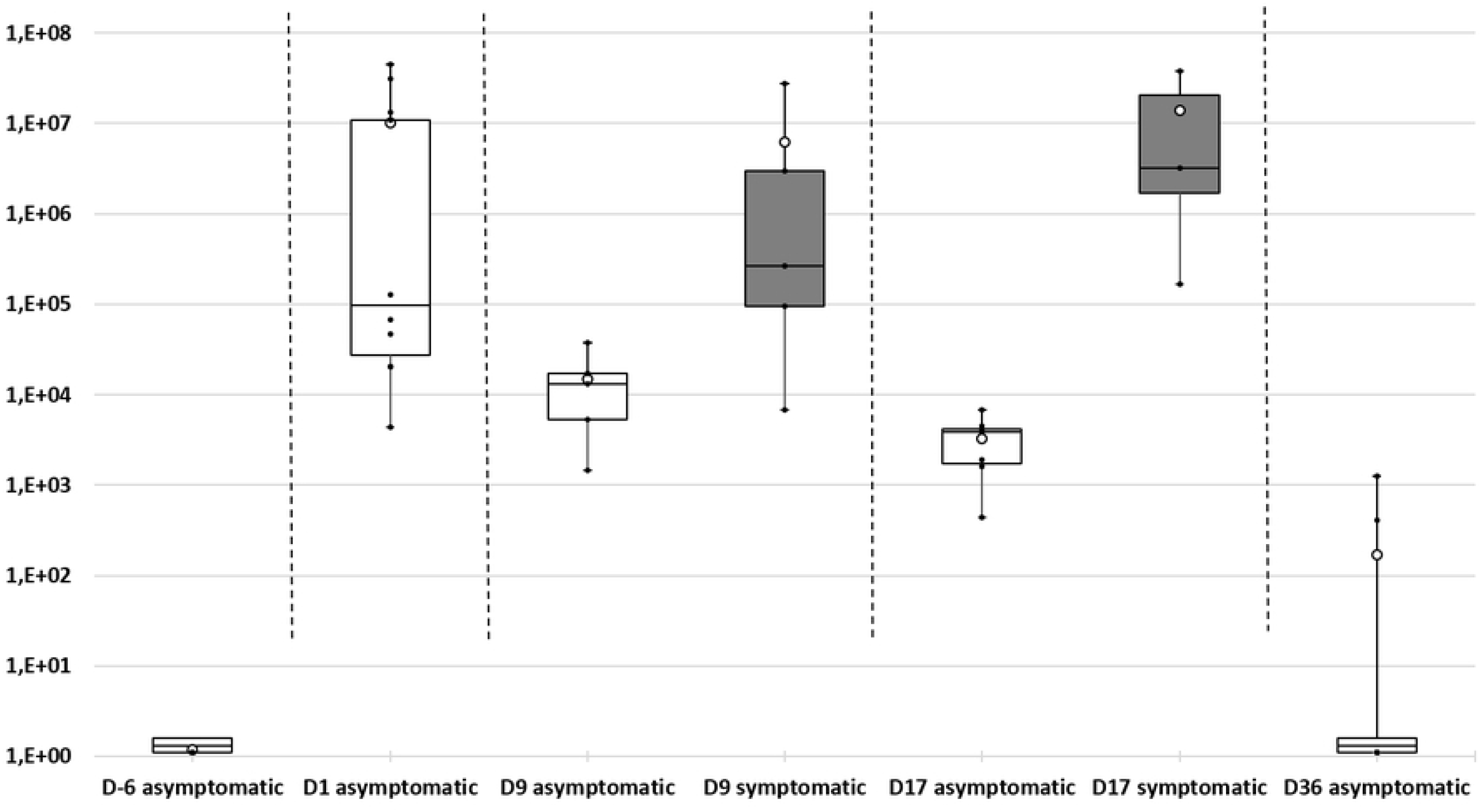
Kinetics of *Tenacibaculum maritimum* bacterial cells quantified by qPCR in the mucus of asymptomatic (white boxes) and symptomatic (grey boxes) batfish from D-6 (at VAIA hatchery) to D36 post transfer to net cage in the lagoon. Quantification results are expressed in numbers of *T. maritimum* cells per μg of total extracted gDNA. Because zero values (no detection) cannot be represented on a logarithmic scale, an arbitrary value of 1.1 was assigned to these negative results. Each box-plot shows mean (white circle), the 25th to 75th percentile (rectangular box), the minimum and maximum values (dots at the extremities), as well as individual quantification (black dots) from 5 to 10 batfish per sampling time and group of batfish.

In contrast, the mucus of all batfish sampled just one day (D1) after the transfer to the lagoon net cages was found positive for *T. maritimum* by qPCR, although very different bacterial loads (mean values, 9.98 10^6^ ± 1.57 10^7^ cells per μg DNA) were observed. At D9, during the first stationary phase of mortality (D6-D13), asymptomatic batfish showed significantly lower bacterial loads in their mucus compared with D1 (Kruskal Wallis test; p = 0.020). Nevertheless, the amount of *T. maritimum* at D9 was significantly higher (p = 0.047) in batfish exhibiting clinical signs of tenacibaculosis compared with asymptomatic fish. During the second peak of mortality, the discrepancy between asymptomatic and symptomatic batfish was even more pronounced (p = 0.016). At the end of the mortality events (D36), *T. maritimum* DNA was absent from most sampled surviving fish (8/10). Indeed, only two asymptomatic batfish among the 10 tested were found positive by qPCR but at very low levels (8.36 10^2^ ± 6.02 10^2^ cells per μg DNA), signalling the end of the outbreak.

qPCR results on the liver of the same batfish sampled for their mucus revealed that only one fish among the 50 tested, sampled at D1, was positive for *T. maritimum* at a low level (1.19 10^4^ cells per μg gDNA). Similar results were obtained with the posterior intestine: only seven batfish were found qPCR positive, with the low value of 1.11 10^4^± 1.37 10^4^ cells per μg gDNA (data not shown).

## 4 Discussion

Orbicular batfish aquaculture in French Polynesia started in 2004 and the first symptoms of white-patch disease were observed in 2006. In this study, we showed that the white-patch disease decimating farmed *Platax orbicularis* is associated with *T. maritimum* infection. To our knowledge, this is the first time that this bacterium has been isolated in French Polynesia, and also the first time it has been retrieved from batfish. *T. maritimum* has been associated with a large variety of marine fish species: 38 according to Nowlan et al. (2020) [5], including the orbicular batfish (this study). Surprisingly, *Platax orbicularis* is the only tropical fish in which this pathogen has been reported so far. Nevertheless, the range of susceptible hosts for this bacterium is probably underestimated. In French Polynesia, tenacibaculosis has dramatic consequences for batfish farms, which often suffer mortality levels over 50%. In addition, the disease may also be of serious concern regarding the diversification of aquaculture programs launched by local authorities.

In this study, we described a reproducible bath challenge protocol that demonstrated that *T. maritimum* is able to infect orbicular batfish by immersion, thus fulfilling Koch’s postulates. Experimental infection procedures using immersion challenges have been broadly used with fish pathogens in recent years because they are likely to mimic the natural infection process more accurately than injection challenges. In particular, immersion does not bypass the first line of fish defence (i.e. the skin mucus barrier), unlike the more common subcutaneous, intraperitoneal and intramuscular injection routes. It has also been reported that, compared with immersion, some injection challenges fail to induce tenacibaculosis [11,26-28] or can lead to high mortality rates in negative controls due to stress and local lesions caused by the injection [29]. However, comparative analyses of challenge protocols are rather difficult to perform due to the many factors reported to influence pathogenicity, such as the bacterial strain [9,27,28], culture conditions (i.e. growth medium and temperature), infection dose [12,26], duration of immersion [30], physical [27] and chemical characteristics of seawater, zootechnical practices (e.g. fish density and animal feed), and the host fish species [8], physiological status (e.g. age [27]) and genetic background (e.g. susceptible, resistant). In this study, no significant difference in mortality rates was observed when batfish were infected with strains TFA4, Aq 16-87, or Aq 16-84, although these strains differed in some genetic traits. However, due to the high virulence of these strains in our immersion challenge model, further studies using a lower dose or shorter immersion time might reveal virulence differences between these strains. Although physical alteration of the fish skin was not necessarily seen before morbidity and mortality, batfish with impaired mucus developed clinical signs more rapidly and experienced higher mortality rates (100% mortality at 23 h post infection) than those which mucus was intact. These results are in agreement with similar studies performed on other fish species [11,30]. Indeed, mucus has been largely documented as an important component of the fish innate immune system and a physical and chemical barrier against pathogens [31].

Some of the isolates from diseased batfish were shown to belong to another *Tenacibaculum* species, *T. mesophilum*, a bacterium initially reported in a marine sponge. In addition, *T. mesophilum* strain HMG1 was shown to degrade malachite green, an antimicrobial that has long been used in aquaculture but now banned in many countries [32]. Immersion challenges performed with strains Aq 9-66 and Aq 9-67 showed that both strains were totally avirulent. Further studies would be needed, however, to determine whether *T. mesophilum* strains can play a role in the pathogenesis of tenacibaculosis, primarily caused by *T. maritimum*, by acting as secondary colonizers of the lesions.

Although this study was conducted with only 10 isolates, an unexpected diversity of *T. maritimum* isolates was found. Our results demonstrate the presence of two distinct groups: strains Aq 16-85, Aq 16-88 and Aq 16-89, belonging to serotype O1 and to sequence type ST167; and strain TFA4, belonging to serotype O3 and to sequence type ST168. Such a diversity among *T. maritimum* isolates was also noticed among the Australian isolates (Fig 4). In agreement with Van Gelderen et al. (2010) [33], no correlation between serotype and geographic distribution was observed in the present study.

Additional work is needed to make a deeper exploration of the genetic diversity of *Tenacibaculum* strains associated with batfish in French Polynesia in order to evaluate their virulence potential and develop management and disease control strategies. Because the natural ecology of *T. maritimum* is still unknown, more in-depth epidemiological studies will also be necessary to decipher the mode of transmission and the natural route of infection of this pathogen.

## Acknowledgements

The authors are indebted to Dr Alicia Estévez Toranzo (Universidad de Santiago de Compostela, Spain) for serotyping the *T. maritimum* batfish isolates. We are also grateful to Sylvain Dupieux and Eddy Laille of Tahiti Fish Aquaculture company, Edouard Lehartel from Tautira Aquaculture company, and Benoît Le Marechal, Director of Cooperative des Aquaculteurs de Polynésie française (CAPF) as well as his colleagues Wallen Teiri, Heifara Wallon and Sylvain Dupieux, who provided us with the experimental fish used in this study. This work was supported by IFREMER and the Direction des Ressources Marines through “Aqua-Sana 1 and Aqua-Sana2” collaborative projects (2016-2021) and by the Ministère de l’Economie Verte et du Domaine en charge des Mines et de la Recherche de Polynésie Française through « Tenacifight » project (2020-2022). P. Lopez was granted an IFREMER-DRM scholarship. We would also like to thank the Bureau de Traduction de l’Université, UBO, Brest, for revising the English of the manuscript.

## Supporting information

**S1 Fig. Phylogenetic relationships of 16S rDNA nucleotide sequences of the strains recovered in this study and the type strains of all *Tenacibaculum* species (See Table S2 for accession number).** The phylogenetic tree was constructed with the MAFFT online tool using the neighbor-joining (NJ) method with Jukes Cantor substitution model on all gap-free sites (1316pb) based on an alignment of 32 members of the genus *Tenacibaculum* performed with the L-INS-i method. Numbers at each branch indicated the percentage bootstrap values on 1,000 replicates. The 16S rDNA sequence of *Pseudotenacibaculum haliotis* (strain FDZSB0410) was used as an outgroup.

**S1 Table.**
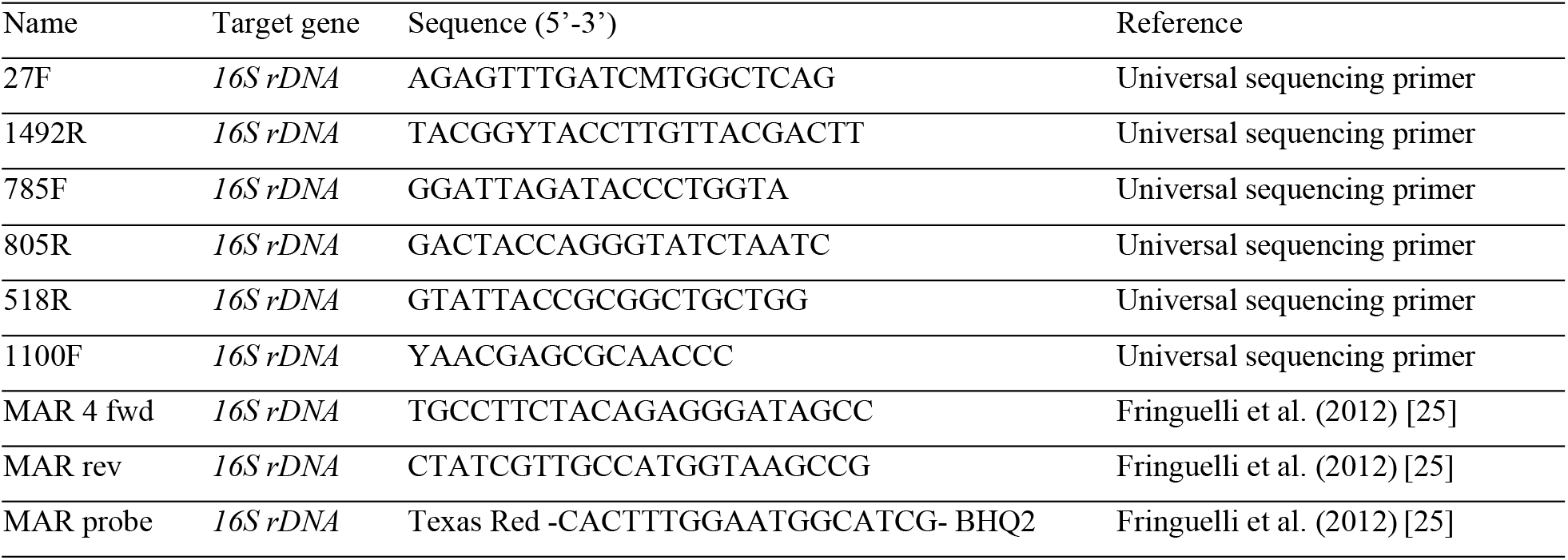
List of PCR primers and probe used in this study.

**S2 Table.** Accession number of the type strains used in S1 Fig.

| Type strains | Genome accession number |
| --- | --- |
| *Tenacibaculum adriaticum* B390 | AM412314 |
| *Tenacibaculum aestuarii* SMK-4 | DQ314760 |
| *Tenacibaculum aestuariivivum* JDTF-79 | MF193601 |
| *Tenacibaculum agarivorans* HZ1 | MSMP01000073 |
| *Tenacibaculum aiptasiae* a4 | EF416572 |
| *Tenacibaculum amylolyticum* MBIC4355 | AB032505 |
| *Tenacibaculum ascidiaceicola* RSS1-6 | KT231981 |
| *Tenacibaculum caenipelagi* HJ-26M | KC832834 |
| *Tenacibaculum crassostreae* JO-1 | EU428783 |
| *Tenacibaculum dicentrarchi* 35/09 | FN545354 |
| *Tenacibaculum discolor* DSM 18842 | RCCS01000002 |
| *Tenacibaculum finnmarkense* DSM 28541 | KT270385 |
| *Tenacibaculum gallaicum* A37.1 | AM746477 |
| *Tenacibaculum geojense* YCS-6 | HQ401023 |
| *Tenacibaculum halocynthiae* P-R2A1-2 | JX912707 |
| *Tenacibaculum haliotis* RA3-2 | KX450476 |
| *Tenacibaculum holothuriorum* S2-2 | LAPZ01000023 |
| *Tenacibaculum insulae* JDTF-31 | MF765760.1 |
| *Tenacibaculum jejuense* KCTC 22618 | LT899436 |
| *Tenacibaculum litopenaei* B-I | DQ822567 |
| *Tenacibaculum litoreum* CL-TF13 | AY962294 |
| *Tenacibaculum lutimaris* DSM 16505 | RAQM01000002 |
| *Tenacibaculum maritimum* NCIMB 2154 | KT270382.1 |
| *Tenacibaculum mesophilum* DSM 13764 | jgi.1107970 |
| *Tenacibaculum ovolyticum* IFO 15947 | AB078058 |
| *Tenacibaculum piscium* TNO020 | GU124766 |
| *Tenacibaculum sediminilitoris* YKTF-3 | KU696540 |
| *Tenacibaculum singaporense* TLL-A2 | MG641897 |
| *Tenacibaculum skagerrakense* D30 | AF469612 |
| *Tenacibaculum soleae* LL04 12.1.7 | AM746476 |
| *Tenacibaculum todarodis* LPB0136 | CP018155 |
| *Tenacibaculum xiamenense* WJ-1 | JX984443 |

